# Direct focus sensing and shaping for high-resolution multi-photon imaging in deep tissue

**DOI:** 10.1101/2021.08.04.455159

**Authors:** Zhongya Qin, Zhentao She, Congping Chen, Wanjie Wu, Jackie K.Y. Lau, Nancy Y. Ip, Jianan Y. Qu

## Abstract

High-resolution optical imaging of deep tissue *in-situ* such as the living brain is fundamentally challenging because of the aberration and scattering of light. In this work, we develop an innovative adaptive optics three-photon microscope based on direct focus sensing and shaping that can accurately measure and effectively compensate for both low- and high-order specimen-induced aberrations and recover near-diffraction-limited performance at depth. A conjugate adaptive optics configuration with remote focusing enables *in vivo* imaging of fine neuronal structures in the mouse cortex through the intact skull up to a depth of 750 µm below pia, making high-resolution microscopy in cortex near non-invasive. Functional calcium imaging with high sensitivity and accuracy, and high-precision laser-mediated microsurgery through the intact skull were demonstrated. Moreover, we also achieved *in vivo* high-resolution imaging of the deep cortex and subcortical hippocampus up to 1.1 mm below pia within the intact brain.

## 1. INTRODUCTION

Breakthroughs in imaging technology have been the primary driving force of new biological discoveries. Optical microscopy has greatly facilitated biomedical research in recent decades, with its ability to provide structural and functional information in living specimens at high spatiotemporal resolution. However, optical aberration and scattering occur as light travels through and interacts with inhomogeneous biological tissues, such as the mammalian brain, fundamentally limiting the performance of optical microscopy in both resolution and depth.

Two-photon microscopy (2PM), the current method of choice for *in vivo* brain imaging of small animals, alleviates the scattering issue by selectively exploiting the ballistic (unscattered) photons for excitation^1^. However, the exponential attenuation of the ballistic photons with imaging depth and the occurrence of out-of-focus fluorescence result in an exponential decrease of the signal-to-background ratio (SBR), limiting the imaging depth to <1 mm for the mammalian brain^2^. Recently, three-photon microscopy (3PM) has shown great potential to extend the imaging depth by using longer wavelength and higher-order nonlinear excitation to reduce the scattering further and suppress the unwanted background fluorescence^3,4^. However, like 2PM, 3PM still suffers from aberrations arising from the variations of refractive index in biological tissues, preventing the ballistic photons from forming a diffraction-limited focus and heavily degrading the spatial resolution^5^. This ultimately prohibits *in vivo* multi-photon imaging being used to resolve subcellular structures in the deep cortical and subcortical layers in a minimally invasive manner.

Adaptive optics (AO) compensation of specimen-induced aberrations is essential to recover optimal imaging performance at depth^6,7^. The desired excitation wavefront can be determined using either direct^8–10^ or indirect^11–14^ approaches. Direct wavefront sensing of a fluorescent guide star inside biological tissue enables rapid measurement of aberration, but it depends on quasi-ballistic photons for wavefront sensing and the performance degrades rapidly at depth^9,10^. In indirect sensing approaches, iterative algorithms are utilized to determine the optimal corrective wavefront. For example, modal AO techniques make use of a deformable mirror that iterates through low-order deformations^11^, and pupil segmentation approaches acquire images through segments of the objective back aperture to estimate the phase gradient in order to correct aberrations^12^. Indirect approaches are more suitable for opaque tissues but are usually time-consuming and can mainly deal with low-order aberrations^11,12^. Recently, an AO technique called focus scanning holographic aberration probing (F-SHARP) has been developed to correct both low- and high-order aberrations at high speed *in vivo* ^15,16^. F-SHARP measures the aberrated electric-field point-spread function (E-field PSF) directly by using two-beam interference, and compensates for the aberrations using phase conjugation of the measured PSF^17^. However, F-SHARP as originally proposed relies on a four-step phase stepping scheme to derive the E-field PSF, which suffers from a poor signal-to-noise ratio (SNR) because the interference signal is weak relative to the strong D.C. background^16^.

In this work, we developed an adaptive optics three-photon microscope (AO-3PM) based on a direct focus sensing and shaping (DFSS) method that shares the basic physical principle as F-SHARP technology. However, DFSS allows accurate measurement and corrective updates of both low- and high-order aberrations in highly scattering tissue at great depth. The method makes use of two innovations: direct focus sensing with phase-sensitive detection and conjugate AO with remote focusing. Specifically, we introduced a high-frequency modulation and phase-sensitive detection scheme to achieve accurate measurement of the E-field PSF in a fast and photon-efficient manner, which allows subsequent aberration correction of a large number of modes by using a high-pixel-count wavefront corrector. In addition, we integrated a remote focusing approach with the conjugate AO configuration to enable effective single correction over a large volume for imaging through a turbid layer such as intact skull of living mouse. Using this conjugate DFSS-AO-3PM technology, we first validated its imaging performance through the intact skull of mouse for both *in vitro* and *in vivo* preparations and achieved near-diffraction-limited spatial resolution with a drastically improved fluorescence signal over a large depth. We then applied it to *in vivo* structural and functional imaging of mouse cortices through the intact skull, and achieved high-resolution imaging up to 750 µm below the pia. Taking advantage of the tight focus provided by DFSS, we demonstrated precise laser microsurgery and investigated the subsequent microglial dynamics in cortex through the intact skull. Further, by using a pupil DFSS-AO-3PM, we achieved high-resolution deep imaging of subcortical structures up to 1.1 mm below pia within the intact brain.

## 2. RESULTS

### 2.1 Conjugate AO-3PM system based on DFSS

Our conjugate DFSS-AO-3PM system consists of three key modules, that is, direct focus sensing, conjugate AO and remote focusing modules, as shown in Fig. 1A and Fig. S1. The system aberrations are corrected by the deformable mirror (DM) using a simple sensorless AO method (see details in *Methods and Materials*). The direct focus sensing in tissue is based on a phase-modulation and lock-in detection scheme (see details in supplementary: *Principle of DFSS*). In order to measure the scattered and aberrated E-field PSF, a weak scanning beam raster scanned over a small field of view (FOV) is introduced to interfere with a strong stationary beam. Thus the three-photon (3P) excited fluorescence signal at scanning coordinate *x* is given by:

**Figure 1.**
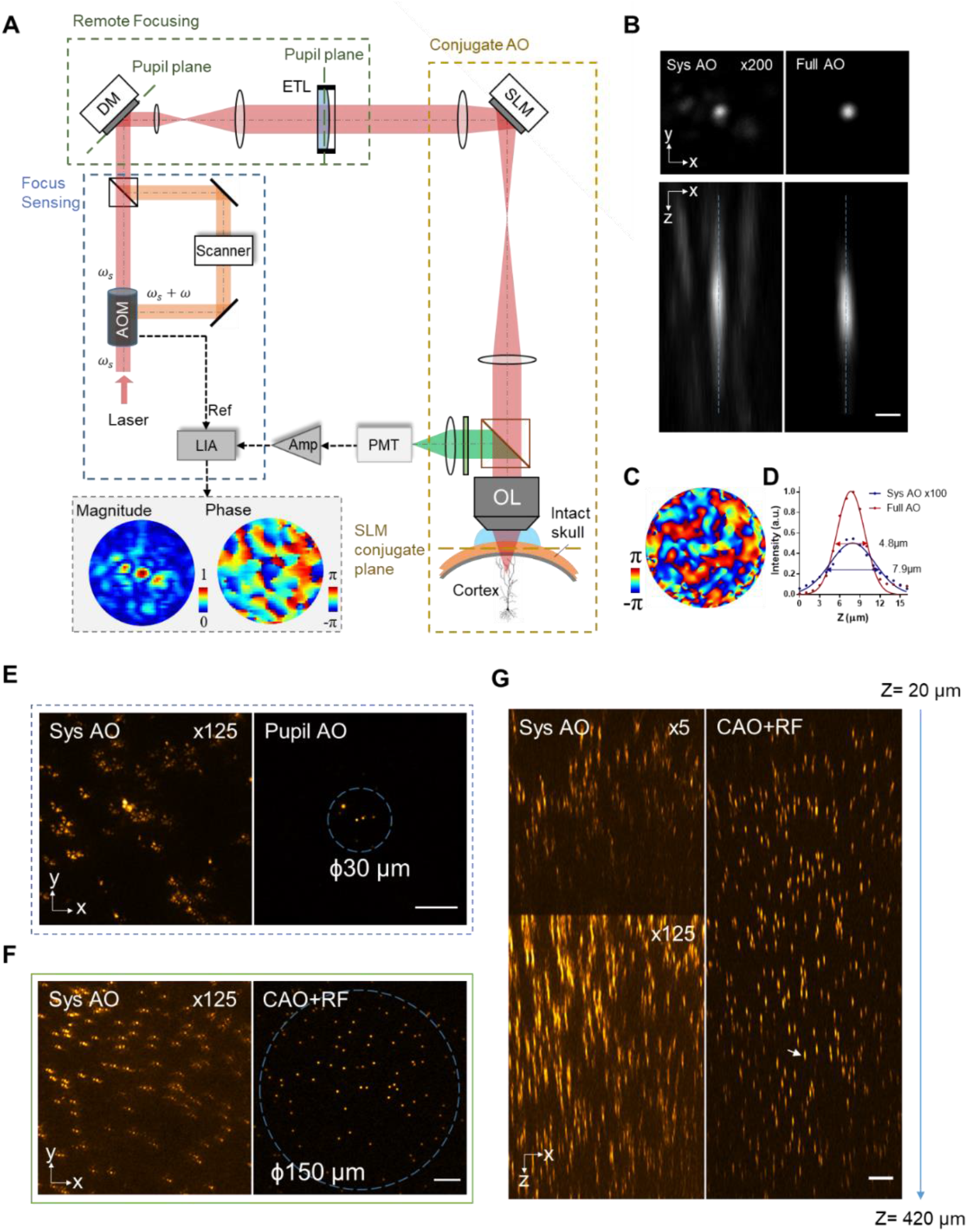
Conjugate DFSS-AO-3PM system and correction of aberrations induced by a 100-µm-thick intact mouse skull. (**A**) Simplified schematic diagram of the conjugate DFSS-AO-3PM system. Blue dashed rectangle: focus sensing module; yellow dashed rectangle: conjugate AO (CAO) module; green dashed rectangle: remote focusing (RF) module. See Fig. S1 for the detailed optical path. (**B**) XY and XZ views of a fluorescent bead (1µm-diameter, 300 µm below skull) image with system correction only (left) and with full correction (right). The images with system correction have been enhanced 200 fold digitally as indicated for better visualization. Scale bar: 2 µm. (**C**) Correction phase pattern applied to the SLM. (**D**) Fluorescence signal profiles along the dashed lines in (B) are plotted with a curve fitted using a Gaussian function. Full width at half maximum (FWHM): 7.9 µm and 4.8 µm for system and full AO, respectively. Note that the theoretical axial resolution for 3PM (1300 nm, 0.7 NA) is ∼3.5 µm. The results showed that near-diffraction-limited resolution was recovered after full AO correction. (**E**) XY view fluorescent beads (300 µm below skull) image with system correction only (left) and with full correction of pupil AO (right). Image of system AO was digitally enhanced as indicated for better visualization. Blue dashed circle indicates the effective FOV. Scale bar: 20 µm. (**F**) XY view of fluorescent beads (300 µm below skull) image with system correction only (left) and with full CAO+RF correction (right). Blue dashed circle indicates the effective FOV. (**G**) XZ view with system correction only (left) and with full CAO+RF correction (right). White arrow indicates the fluorescent bead used for aberration measurement. Scale bar: 20 µm.

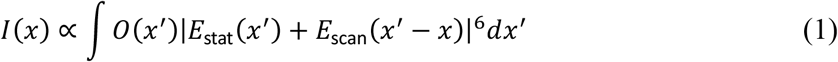

Where *E*_stat_ and *E*_scan_ are the complex-valued E-field PSF of the stationary and scanning beam, and *O*(*x*′)is the real-valued object function related to the fluorophore distribution in the focal plane. By modulating the phase of scanning beam at a fixed frequency *ω* and setting the scanning beam to be much weaker than the stationary beam (i.e., |*E*_scan_|^2^/|*E*_stat_|^2^ < 0.1), the 3P excited fluorescence becomes

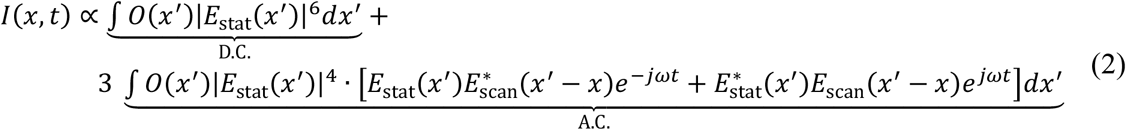

The fluorescence signal *I*(*x, t*)was then demodulated using a lock-in amplifier at frequency *ω*, outputting the orthogonal *X* (i.e., in-phase) and *Y* (i.e. quadrature) as

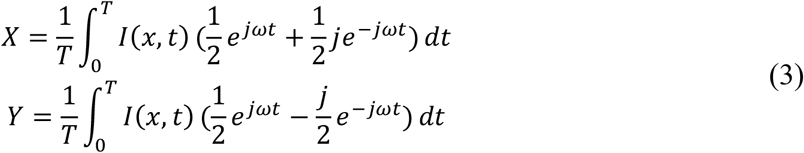

We may then compute the E-field PSF as

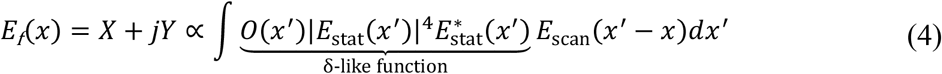

Owing to the high sensitivity provided by our phase-modulation and lock-in detection, the E-field PSF can be determined reliably by scanning a single frame of the FOV, regardless of the large D.C. background. By measuring the aberrated E-field PSF, the desired corrective wavefront can be derived by using phase conjugation of the measured PSF. Also, a high-pixel-count spatial light modulator (SLM) acting as the wavefront corrector is then employed to correct the aberrations over a large number of modes^15,16^.

To evaluate the efficacy of our DFSS-AO-3PM and characterize its performance in through-skull imaging, we first conducted an *in vitro* experiment by imaging fluorescent beads embedded in agarose below an excised intact skull. In order to measure the wavefront distortions, the strong beam was parked at the brightest regions, e.g. the center of fluorescent bead, while the weak beam was used to scan over a 16×16 µm^2^ FOV to capture the complex E-field PSF. Thanks to the phase-sensitive detection scheme, the scattered and aberrated E-field PSF through the intact skull can be obtained accurately, as shown in Fig. 1A. The overall aberrations can then be accurately determined and effectively corrected by using the two-dimensional Fourier transform of the measured E-field PSF, in a fast manner with only 2-3 iterations (Fig. S2). As can been seen, the fluorescent bead image recorded with system aberration correction only is highly distorted, and the appearance of the side lobes indicates the high-order aberrations induced by the thick skull (Fig. 1B). In contrast, by applying the corrective phase pattern to the SLM to compensate for specimen-induced aberrations (Fig. 1C), we effectively recovered the near-diffraction-limited resolution, and improved the fluorescence intensity more than 200 fold (Fig. 1B and D). These results demonstrated that the DFSS-AO approach enables accurate determination and effective correction of aberrations in 3P imaging through the intact skull.

In AO microscope systems, the wavefront corrector is usually projected to the rear pupil plane of the objective (Fig. S3A). This convenient configuration, known as pupil AO, is implemented in almost all AO microscopes. However, pupil AO correction can only restore the focus quality over a very small FOV (< 30 µm in diameter), as shown in Fig. 1E, because of the rapid decorrelation between the scanned corrective wavefront and the turbid skull with a highly heterogeneous distribution of refractive index. Since the mouse skull is the dominant aberration layer and accounts for most of the deterioration in transcranial brain imaging at depth, the effective FOV after AO correction can in principle be enlarged by conjugating the wavefront corrector to the skull layer, instead of the objective pupil plane. This approach, called conjugate AO, has been demonstrated to increase the corrective FOV in transcranial brain imaging *in vivo*^16,18^. However, these previous implementations of conjugate AO still have a limited axial FOV. To achieve a large effective imaging volume with a single AO correction, we integrated conjugate AO with remote focusing to maintain the conjugation between wavefront corrector and skull layer during axial scanning using an electrical tunable lens (ETL). Here, the aberrations induced by remote focusing are counted as part of system aberrations and corrected by the deformable mirror prior to DFSS-AO corrections (see details in *Method and Materials*). As can be seen, the effective FOV after a single correction of conjugate AO is extended to 150 µm and 400 µm in the lateral and axial directions (Fig. 1F-G and Fig. S3C-D), respectively, which corresponds to an increase of corrective volume by more than 100 fold compared with pupil AO configuration. Moreover, because axial scanning is achieved through remote focusing without requiring physical translation of the objective and wavefront corrector, fast axial scanning across different imaging planes can be easily achieved. Indeed, it should be noted that the enhancement of fluorescence intensity decreases when the imaging plane is away from the correction depth at 300 µm (Fig. S3C-D). This may be due to the fact that the intact skull (100µm thick) is not an infinitely thin layer of dominant aberration and the corrective phase pattern measured at one location is not optimal for other locations. In addition, for the imaging at depth > 300 µm, the excitation beam on SLM is larger than the corrective phase pattern, resulting in incomplete correction for the aberrations and reduced effective NA.

We next validate the ability of conjugate DFSS-AO-3PM to improve *in vivo* brain imaging through the intact skull. We obtained *in vivo* images of microglia labeled with green fluorescent protein (GFP) in adult Cx3Cr1-GFP mice in either pupil or conjugate AO configuration. As shown in Fig. S4, the detailed structures of microglia at 300 µm below the intact skull can barely be identified using a conventional 3P microscope (system AO correction). By using direct focus sensing with the GFP signal, pupil AO effectively compensates for the specimen-induced aberrations and successfully recovers the processes of microglia with less than 3 iterations (Fig. S5), but its effective FOV is restricted to less than 30 µm (Fig. S4A). By contrast, conjugate AO not only delivers similar improvements in resolution and signal intensity as those of pupil AO, enabling microglial processes to be clearly resolved, but also significantly enlarges the corrected FOV to 120 µm (Fig. S4B and F). Furthermore, conjugate AO with remote focusing enables effective improvement of imaging resolution over large imaging depths ranged from 100 to 500 µm with a single corrective wavefront at 300 µm (Fig. S6 and Movie S1). It was found that despite that the aberrations in deeper brain regions (e.g. at 490 µm) may require a corrective phase pattern covering a larger skull area, the corrective wavefront measured at 300 µm still shows an improved signal intensity (3-fold), as shown in Fig. S7. This signal gain is essential for subsequent direct focus sensing at 490 µm due to the limited photon budget at depth. Therefore, we developed a pre-compensation strategy to improve imaging in deeper regions of the brain. Specifically, the additional corrective phase measured at current depth was added to that measured at the previous depth to generate the final and optimal phase pattern for full AO correction (Fig. S7C). In this way, we can effectively measure the aberrations at depth and further enhance the signal intensity and spatial resolution (Fig. S7A-B). These results reveal the great advantages of conjugate DFSS-AO with remote focusing for *in vivo* high-resolution imaging of mouse cortices through the intact skull over large volumes.

### 2.2 High-resolution imaging of brain structure and function through the intact skull

Having demonstrated the ability of conjugate DFSS-AO to improve through-skull imaging, we next conducted *in vivo* imaging of neuronal and vascular structures in the mouse cerebral cortex through the intact skull. Fig. 2A shows a cortical column of an adult Thy1-YFP transgenic mouse with a 100µm-thick skull. Detailed experimental parameters are summarized in Table S1. Here, dual-color fluorescence imaging of YFP-labeled pyramidal neurons and Texas-Red-labeled cerebral microvasculature is achieved with single-wavelength (1300 nm) 3P excitation^19^. As can be seen, even with system aberrations corrected, contrast and resolution in 3P imaging through the intact skull degrades rapidly with increase of depth (Fig. 2A and Fig. S8), primarily because of the distortion of excitation PSF caused by skull-induced aberrations. The deterioration in PSF is particularly problematic in 3P imaging because the fluorescence has a third-order dependence on excitation density^3^. Thus small neuronal dendrites beyond 200 µm depth can barely be resolved by conventional 3P imaging through the intact skull ^20^ (Fig. S8). With the highly sensitive DFSS method, both the low- and high-order aberrations experienced by the excitation light can be determined accurately and corrected effectively with 2-3 iterations. In this way, AO recovery of near-diffraction-limited PSF leads to much improved resolution with significantly stronger signals (up to 30 fold), allowing fine structures such as neuronal dendrites and individual spines to be visualized clearly through the intact skull (Fig. 2B-I and Fig. S8). Through-skull imaging of microvascular structures is also significantly improved after full AO correction, with much increased SBR at depths (Fig. S9). Quantitative measurement of the fluorescence intensity profiles of dendrites at depth provides an upper-bound estimation of the axial resolutions (Fig. S10), indicating that conjugate DFSS-AO-3P successfully restores the near-diffraction-limited resolution up to 750 µm below pia.

**Figure 2.**
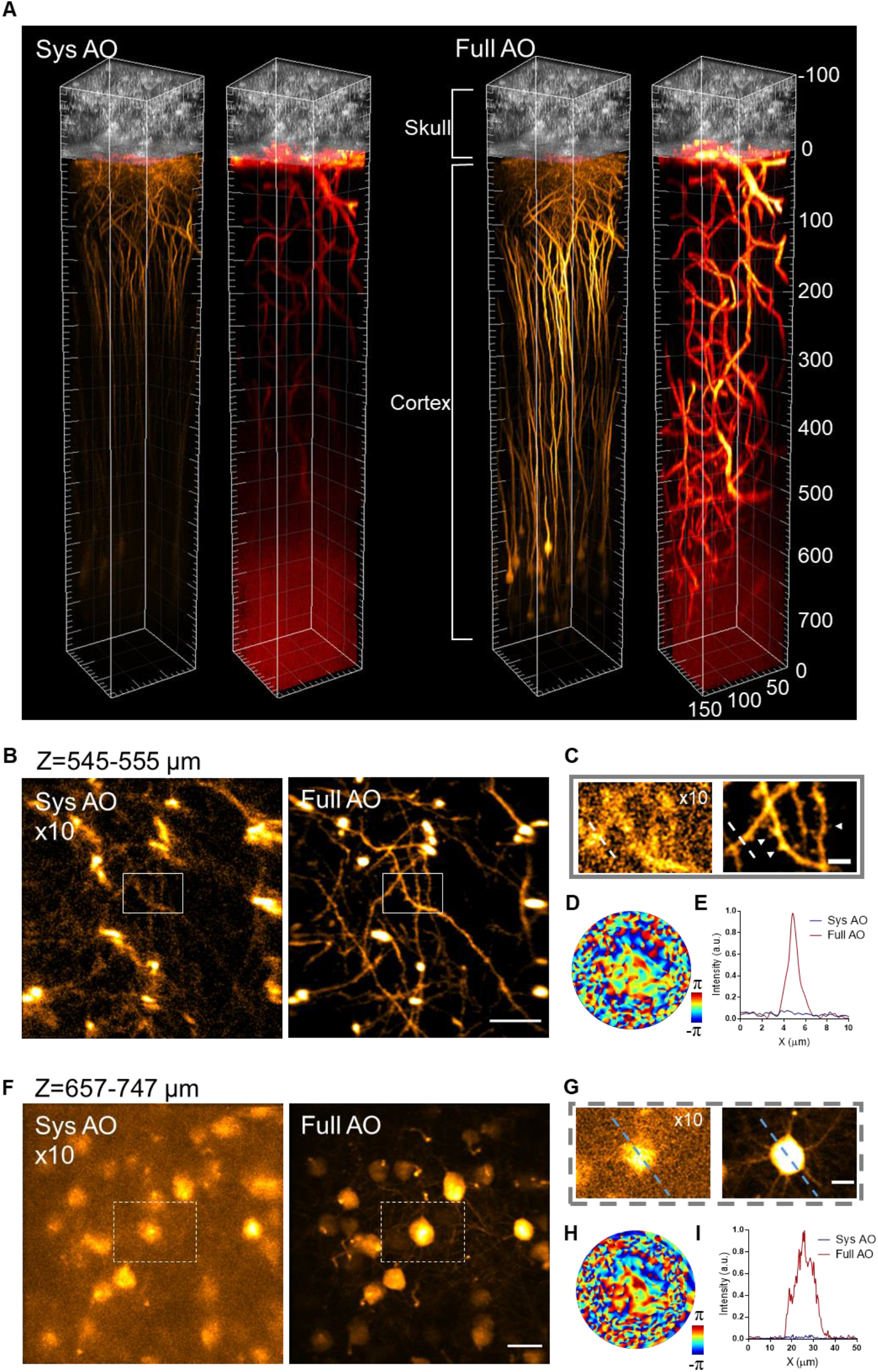
Conjugate DFSS-AO-3PM enables *in vivo* cortical imaging with near-diffraction-limited resolution over large volumes through the intact skull. (**A**) 3D reconstructions of *in vivo* dual-color 3P imaging of YFP-labeled neurons (orange) and Texas Red Dextran-labeled blood vessels (red) in a 150×150×780µm^3^ cortical column of a 2-month-old Thy1-YFP transgenic mouse through 100-µm-thickness intact skull. Signal from skull (grey) is THG. The zero depth is set just beneath the pia. (**B**) Maximum intensity projection (MIP) of stack images (545-555 µm below pia) of YFP-labeled pyramidal neurons from (A) with system correction only (left) and with full correction (right). The image of system AO correction was digitally enhanced as indicated for better visualization. Scale bar: 20 µm. (**C**) Magnified views of dendrites in white box in (B). White arrow heads indicate the dendritic spines. Scale bar: 5 µm. (**D**) Correction phase pattern of SLM for the full AO image in (B). (**E**) Signal profiles along the dashed line in (C). (**F**) MIP of stack images (657-747 µm below pia) of pyramidal neurons from (A) with only system correction (left) and with full correction (right). Scale bar: 20 µm. (**G**) Magnified views of neurons in white box in (F). Scale bar: 10 µm. (**H**) Correction phase pattern of SLM for the full AO image in (F). (**I**) Signal profiles along the dashed line in (G).

By virtue of the rapid direct focus sensing, we also applied the conjugate DFSS-AO-3PM to functional calcium imaging of neuronal activity, in which the fluorescence levels change rapidly and cannot be used for aberration measurement in most indirect AO approaches^21,22^. For direct focus sensing to work effectively, we normalized the outputs of lock-in detected signal (A.C. term) to the fluorescence signal (D.C. term) during focus sensing with fluctuating fluorescence (see details in *Methods and Materials*). In this way, the artifact in the measured E-field PSF caused by rapidly varying calcium fluorescence can be eliminated, enabling the aberrations to be determined accurately (Fig. S11). We next performed *in vivo* calcium imaging of GCaMP6s-labeled neurons in the somatosensory cortex of CCK-GCaMP6s transgenic mice through the intact skull. As shown in Fig. 3, the spontaneous activity of neurons became barely discernable at depth for conventional 3P imaging (system AO correction) because of the overwhelming background noise or so called neuropil contaminations ^23^, which are especially problematic in the densely-labeled brain. By using our AO approach, the fluorescence signals from the neuronal somata and dendrites were increased significantly together with drastically improved resolution, enabling the spontaneous neuronal activity to be recorded accurately at over 445 µm below the cortical surface. In addition, due to the large imaging volume and fast axial scanning provided by the conjugate AO with remote focusing method, we demonstrated near-simultaneous multi-plane calcium imaging of neuronal and dendritic activities from different cortical layers through the intact skull (Fig. S12 and Movie S2).

**Figure 3.**
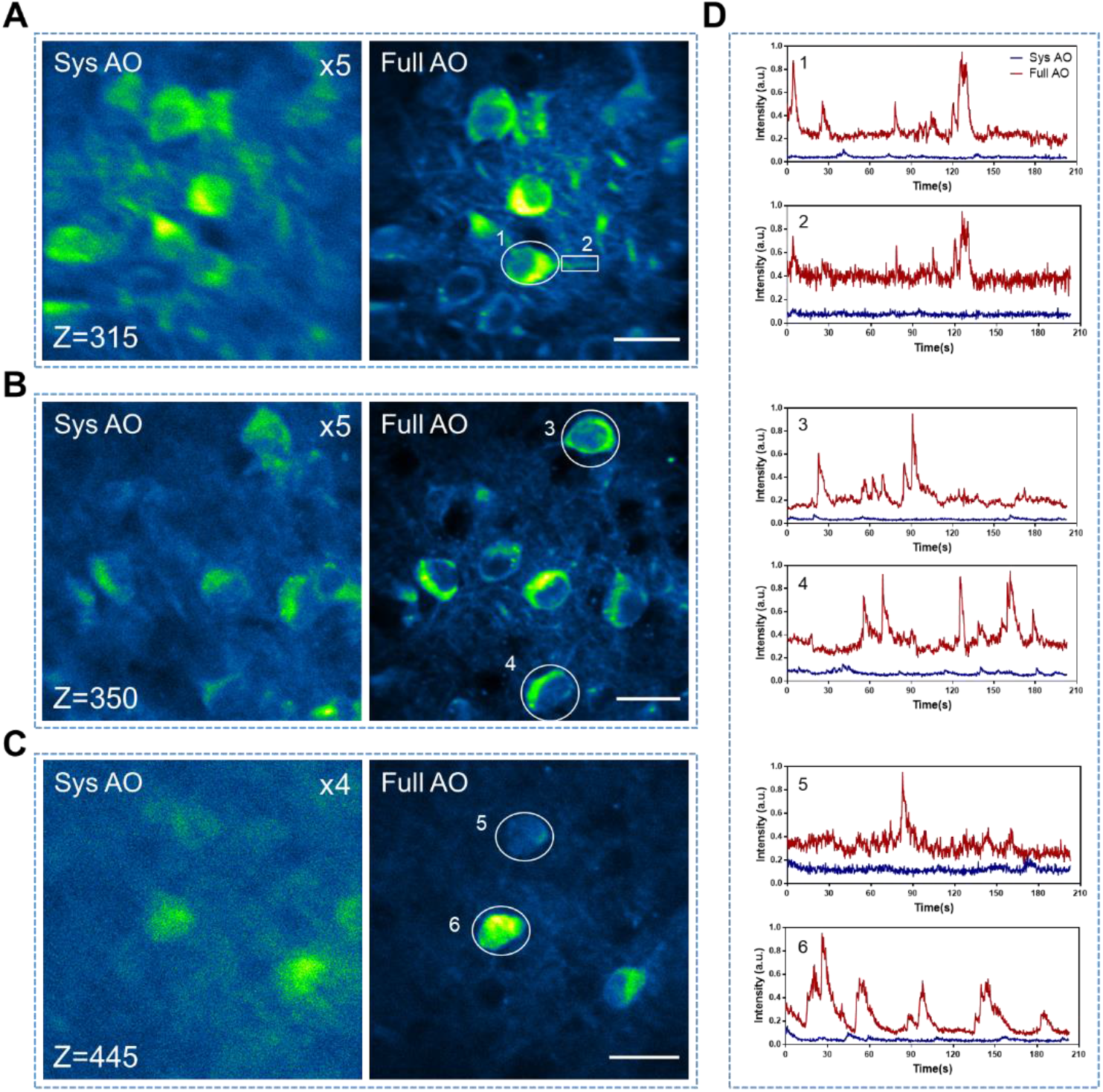
Conjugate DFSS-AO-3PM improves *in vivo* functional calcium imaging of neuronal activity through the intact skull. (**A**-**C**) *In vivo* 3P calcium imaging of spontaneous neuronal activity at different depths below pia of the somatosensory cortex of a CCK-GCaMP6s mouse (4-month-old) through the intact skull (110µm-thickness) with system correction only (left) and full AO correction (right). Images are acquired at 4.43 fps and shown as average-intensity projections of 900 frames. Scale bar: 20 µm. (**D**) Fluorescence traces extracted from the numbered ROIs as shown in (A-C).

Microglia, the primary resident immune cells of the central nervous system, play essential roles in brain homeostasis and neurodegenerative diseases^24^. Noninvasive imaging tools with high-resolution capability are crucial for the study of undisturbed microglial physiology. Taking advantage of our conjugate DFSS-AO approach, we next conducted *in vivo* through-skull imaging of microglia in the cerebral cortex of Cx3Cr1-GFP transgenic mice. As can be seen in Fig. 4A-C, conjugate AO consistently improved the image resolution and fluorescence signal, enabling us to resolve the fine processes of resting ramified microglia over large imaging volumes, which were otherwise not discernable without full AO correction. Moreover, by compensating for the specimen-induced aberrations and recovering a tight laser focus inside the cortex, DFSS allows us to perform precise laser microsurgery through the intact skull, which is a powerful technique for studying the cellular mechanisms underlying various pathological conditions ^25,26^. Time-lapse imaging at multiple depths revealed that the highly-localized lesion only activated a few adjacent microglia (within a distance of 50 µm), which rapidly extended their processes towards the lesion in a coordinated manner (Fig. 4A and Movie. S3). These results revealed the great potential of DFSS for precise optical manipulation, in additional to high-resolution imaging of brain through the intact skull.

**Figure 4.**
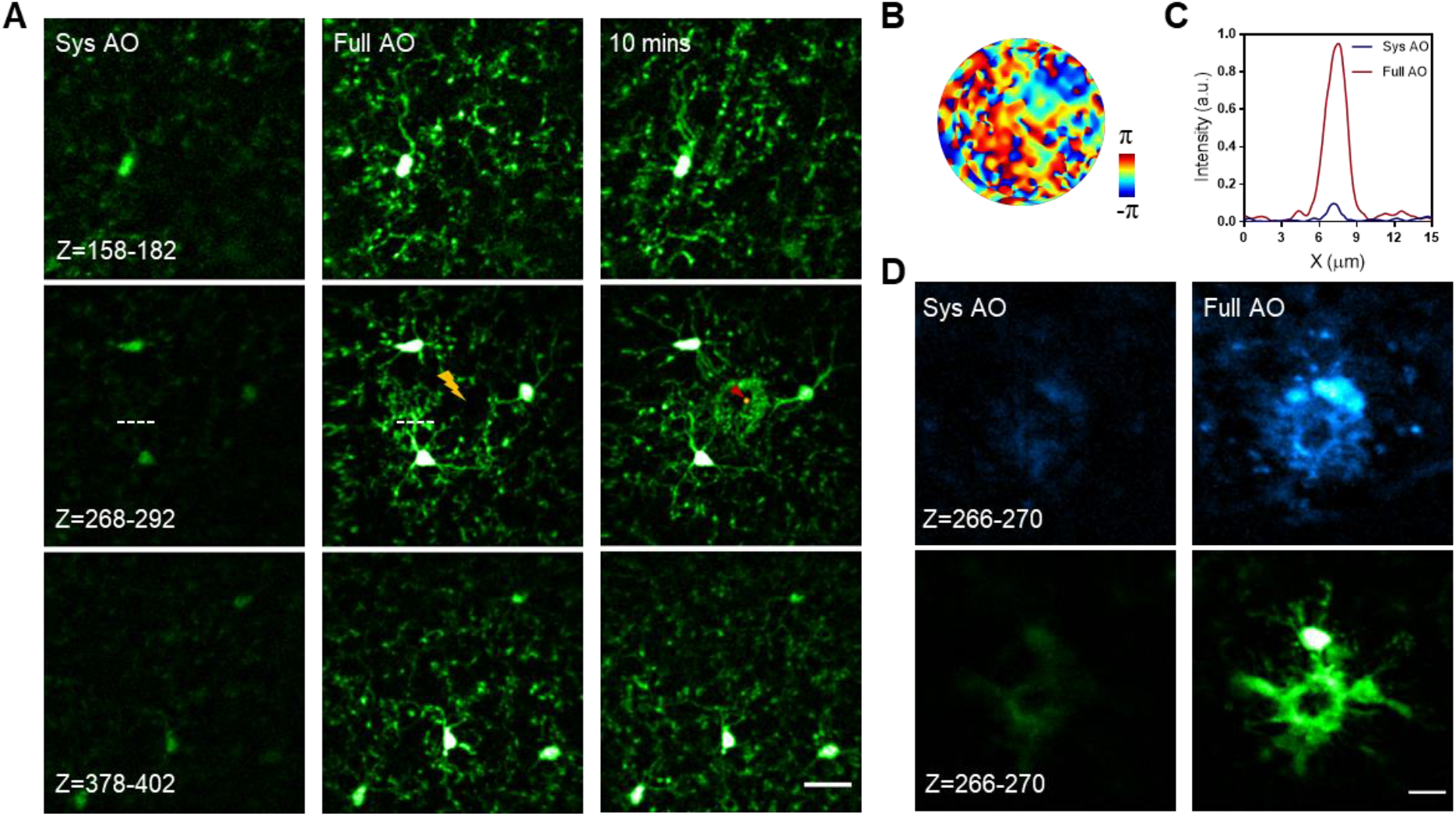
Conjugate DFSS-AO-3PM enables precise laser micro-lesion and *in vivo* high-resolution imaging of microglia in aged AD brain through the intact skull. (**A**) Time-lapse images of GFP-labeled microglia in Cx3Cr1-GFP transgenic mouse (2-month-old) following precise laser micro-lesion. Images are shown as MIP acquired at different imaging depths with system correction only (left) and with full correction (middle and right). The lightning symbols indicates the spot for subsequent laser ablation. The red arrow in the 10-mins image indicates the localized spot of micro-lesion. Scale bar: 20 µm. (**B**) Correction phase pattern of SLM for the full AO image in (A). (**C**) Signal profiles along the dashed line in (A). (**D**) Dual-color imaging of microglia expressing GFP (green channel) and amyloid plaque labeled by MeO-X04 (blue channel) in a 12-month-old APP/PS1×Cx3Cr1-GFP mouse through a 140µm-thick intact skull. Scale bar: 10 µm.

To further assess the capability of our conjugate DFSS-AO in improving through-skull imaging, we then conducted *in vivo* cortical imaging of aged (>10-month-old) mice with thicker skulls (130-140 µm). Due to the severe scattering and aberration induced by the irregular skull in aged mice, the conventional 3P imaging (system AO) is unable to produce reasonable resolution at depths over 100 µm below pia (Fig. S13). With DFSS to correct for the aberrations, however, the imaging resolution was recovered to near-diffraction-limited level and the fluorescence signal was substantially improved, enabling dendritic spines to be visualized clearly. The complex pattern of corrective phase indicates that DFSS compensates the higher-order aberrations to achieve high-resolution imaging through the thicker intact skull of aged mice. By using conjugate AO, we successfully maintained high-resolution 3P imaging over 500 µm below pia. In addition, we also performed *in vivo* through-skull imaging of microglia in a mouse model of Alzheimer’s disease (AD), in which amyloid plaques accumulate in the brain through aging and are thought to be the causative agent of AD pathology^27^. Owing to the high resolution provided by DFSS, we can clearly resolve the morphology of microglia surrounding the amyloid plaques through a 140µm-thick intact skull which were otherwise unidentifiable without full AO correction (Fig. 4D). The observed morphology of the plaque-associated microglia is consistent with our previous study^28,29^. These results demonstrate that our AO-assisted through-skull imaging approach can facilitate *in vivo* studies of aged mouse brain.

We have demonstrated that conjugate DFSS-AO-3PM is an ideal near non-invasive approach for brain imaging. On the other hand, the skull can also be thinned mechanically down to a certain thickness (∼50 µm) to provide a minimally invasive access to the brain with reduced scattering^30^. Despite conventional 3P imaging achieving a larger penetration depth through thinned skull than through intact skull, its spatial resolution still degrades rapidly at depth, being insufficient to identify tiny neuronal structures such as dendritic spines beyond 400 µm below the pia (Fig. S14). We further demonstrated that our conjugate DFSS-AO approach also significantly improved *in vivo* cortical imaging through a thinned skull window, recovering near-diffraction-limited resolution over a large depth (>700 µm) similar to that of intact skull (Fig. S14 and Fig. S15). Moreover, functional calcium imaging through a thinned skull window is also drastically improved after full AO correction (Fig. S16), and we achieved highly-sensitive and accurate recording of spontaneous calcium activity from neuronal somata and dendrites on different planes up to 600µm below the cortical surface (Fig. S17).

### 2.3 High-resolution imaging of deep brain through an open skull window

To push the imaging depth further, we performed 3P imaging through an open skull window, where a section of skull bone is replaced by a transparent glass to provide direct optical access to the brain ^31,32^. In this case, the brain remains intact and specimen-induced aberration mainly arises from the brain tissue which accumulates with imaging depth, without contribution from any specific dominant layer. Therefore, we reconfigured our AO-3P microscope system into the pupil conjugation with a high excitation NA of 1.05 to image the mouse brain at a higher resolution (see *Methods and Materials*). We imaged the pyramidal neurons throughout the cerebral cortex *in vivo*, as shown in Fig. 5A. Here, DFSS leads to significant increases in 3P signal (∼9-fold) and recovery of near-diffraction-limited resolution in the deepest cortical layer, allowing individual spines of basal dendrites to be clearly visualized (Fig. 5B-D and Fig. S18).

**Figure 5.**
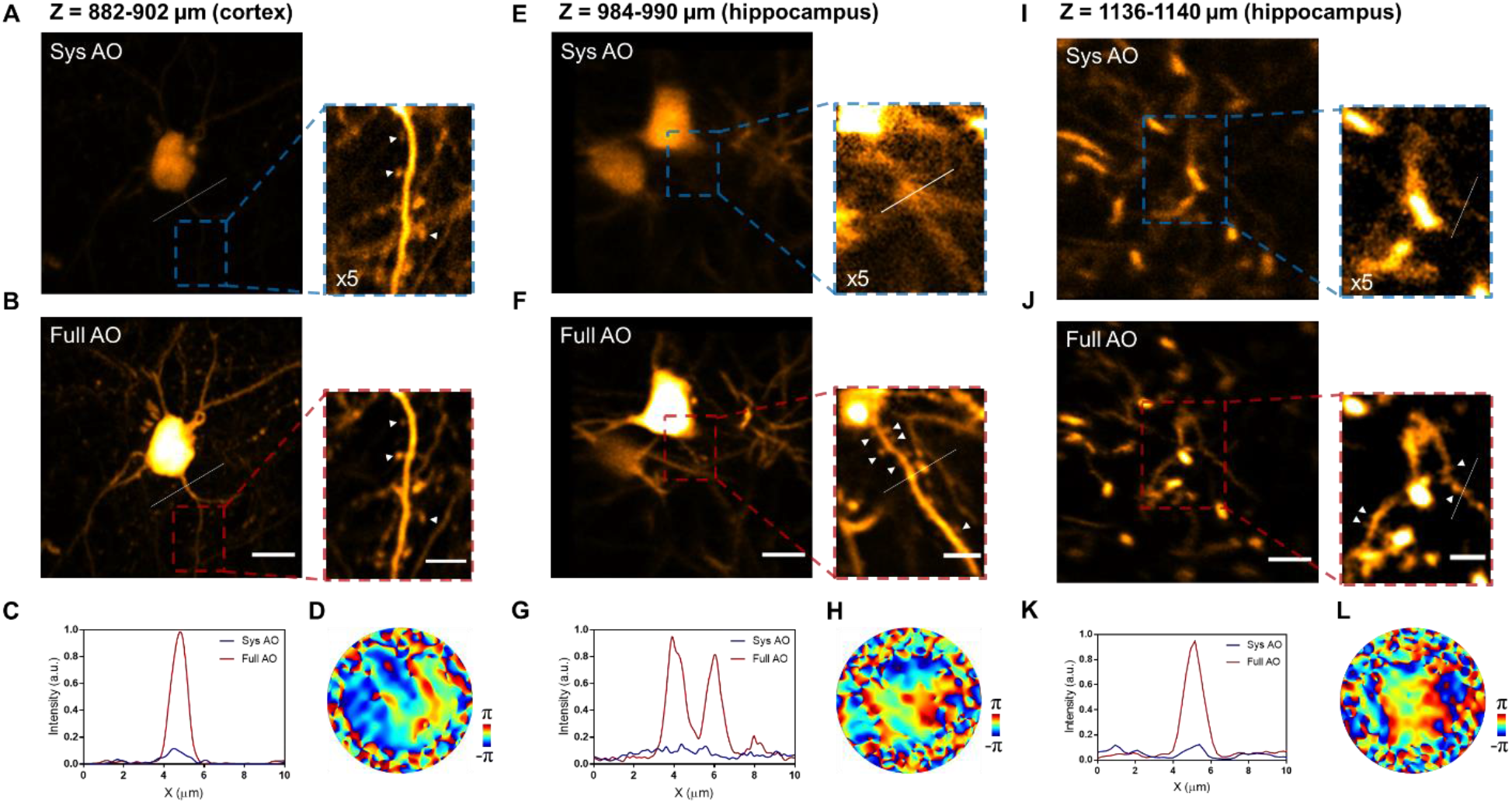
Pupil DFSS-AO-3PM enables *in vivo* imaging of deep-cortical and hippocampal neurons at synaptic resolution through an open skull window. (**A**-**B**) MIP images of GFP-labeled cortical neurons (Z=882-902 µm below pia) in Thy1-GFP mouse (2-month-old) with system correction only (A) and with full correction (B). Scale bar: 10 µm. Insets display magnified views of neuronal dendrites from the outlined regions. White arrow heads indicate the dendritic spines. Scale bar: 5 µm. (**C**) Signal profiles along the white line in (A-B). (**D**) Correction phase pattern of SLM in (B). (**E**-**F**) MIP images of hippocampal CA1 neuron (Z=984-990 µm below pia) in Thy1-GFP mouse with system correction only (E) and with full correction (F). Scale bar: 10 µm and 5 µm (insets). (**G**) Signal profiles along the white line in insets of (E-F). (**H**) Correction phase pattern of SLM in (F). (**I**-**J**) MIP images of hippocampal CA1 neuron (Z=1136-1140 µm below pia) with system correction only (I) and with full correction (J). Scale bar: 10 µm and 5 µm (insets). (**K**) Signal profiles along the white line in insets of (I-J). (**L**) Correction phase pattern of SLM in (J).

Because of limits imposed by the thick and scattering cerebral cortex, it still remains a challenging task to achieve *in vivo* high-resolution imaging beyond the cortex and into the subcortical regions (e.g. hippocampus) without using highly invasive procedures such as GRIN lens insertion^33,34^ and cortical tissue aspiration^35^. Previous work on 3P imaging of the hippocampus *in vivo* provided imaging resolution at the cellular level ^3,4^. As shown in Fig. 5E, despite the neuronal somata in hippocampus CA1 being identifiable in conventional 3P imaging (system AO), the dendritic structures is still unresolvable due to brain-induced aberration (Fig. 5E and I). With pupil DFSS-AO, however, we substantially improve the 3P fluorescence signal and successfully restored near-diffraction-limited resolution, allowing the apical dendritic spines of hippocampal CA1 neurons to be clearly resolved at depths up to 1140 µm below pia (Fig.5 E-L and Fig. S19).

## 3. DISCUSSION

The performance of optical microscopy is greatly affected by scattering and aberration arising from the optical heterogeneity of biological tissues, which rapidly degrade the imaging resolution at depth. In this study, we have developed an adaptive optics three-photon microscopy technique that incorporates two innovations: direct focus sensing with a phase-sensitive detection and conjugate AO with remote focusing. The phase-sensitive detection scheme is crucial for the accurate determination of the scattered and aberrated focal field, thus allowing effective correction of both low- and high-order aberrations induced by the specimen. Conjugate AO integrated with remote focusing ensures that a single corrective phase pattern can reliably recover near-diffraction-limited resolution over a large imaging volume below a single turbid layer. Using the conjugate DFSS-AO-3PM system, we performed *in vivo* brain imaging through the intact skull, a desirable near-noninvasive but also considerably challenging approach. By correcting the specimen-induced aberrations, we achieved *in vivo* high-resolution imaging of mouse cortices up to 750 µm below the pia through the intact skull. With the great speed of direct focus sensing, we showed *in vivo* functional calcium imaging of densely-labeled neurons at depth with drastically improved sensitivity and accuracy. By virtue of the tight focus recovered by AO, we also demonstrated precise laser microsurgery in mouse cortices through the intact skull, and high-resolution dynamic imaging of the microglial responses. To push the depth limit further, we also performed *in vivo* imaging through the open skull window with pupil AO, which enables *in vivo* imaging of deep-cortical and subcortical structures at synaptic resolution up to 1.1 mm below pia within the intact brain. Our results demonstrate that DFSS-AO-3PM technology holds great potential to advance *in vivo* imaging techniques and facilitate neuroscience studies of the living brain.

In this work, we showed that direct focus sensing with phase-sensitive detection enables us to accurately reconstruct the E-field PSF by using the fluorescence signal from either the stable cellular markers (e.g. GFP) or the fluctuating functional reporters (e.g. GCaMP6s). It should be noted that direct focus sensing can also be achieved by either fluorescence or THG signals from blood vessel, although the accuracy is slightly compromised by artefacts arising from the motion of red blood cells (Fig. S20). Though we have demonstrated the capability of DFSS-AO-3PM for deep brain imaging at 1300 nm excitation, it can be applied to 1700 nm excitation without any difficulty. Theoretically, for a larger imaging depth, 3P excitation at 1700 nm is advantageous, due to the longer effective attenuation length (∼ 300 µm for 1300 nm and ∼ 400 µm for 1700 nm in mouse grey matter)^3^. However, the imaging depth will be still limited by the signal depletion in deeper brain due to the inefficient 3P excitation. It has been reported that quantum dots have a 3P excitation cross section of four to five orders of magnitude larger than those of conventional fluorescent dyes and have enabled 3P imaging of mouse cerebral vasculature at up to 2.1 mm depth^36^. As the 3P fluorescence signal from blood vessel can also be used for DFSS, it may serve as a start point for full correction of aberrations in deeper brain region, which first drastically boost the fluorescence intensity of cellular structures labeled with fluorescent markers and allows for a second correction of residual aberrations based on the fluorescent proteins.

## 4. METHODS AND MATERIALS

### Conjugate DFSS-AO-3PM system

A schematic diagram of our 3P microscope system is shown in Fig. S1. The excitation laser (1300 nm, 1 MHz repetition rate) was from a two-stage optical parametric amplifier (Orpheus-F, Light Conversion) pumped by a 40 W femtosecond laser (Carbide, Light Conversion). A homebuilt single-prism pulse compressor was used to compensate for the group delay dispersion of the microscope system and laser source, yielding a final pulse duration of 60fs reaching the sample^37^. The excitation laser then passed through an acousto-optic modulator (AOMO 3080-122, Crystal Technology) driven by a RF signal generator (AODS20160-8, Crystal Technology). The 0^th^ and 1^st^ order diffracted beams were used as the stationary and scanning beams, respectively. The scanning beam was expanded to match the aperture of a XY galvanometric scanning mirror (GVS002, Thorlabs). An optical delay line (not shown in Fig. S1) was placed in the scanning beam arm to match the path length of the two interferometer arms. The two beams were then combined by a polarizing beamsplitter and expanded to slightly overfill the aperture of the DM (DM97-15, Alpao), which was used to correct for system aberrations based a simple sensorless method^38^. Because the two beams have orthogonal polarizations, a half wave plate and a polarizer were placed before the DM to ensure the same polarization and also to adjust the intensity ratio of these two beams. The DM was conjugated to the 3-mm x-galvanometric scanning mirror (6215H, Cambridge Technology) via a 4*f* system (L7: f = 230 mm, consisted of LA1484-C and LA1464-C; L8: f=50 mm, AC127-050-C, Thorlabs). The x and y scanning mirrors were mutually conjugated by lens pair L9 and L10, each of which consisted of two doublets (f=100 mm, #45-806, Edmund Optics) in Ploessl configuration. Then the galvanometric y-scanning mirror was conjugated to an ETL (EL-16-40-TC-NIR-5D-C, Optotune) by scan lens L11 (f=50 mm, SL50-2P2, Thorlabs) and tube lens L12 (f=250 mm, AC508-250-C, Thorlabs). Last, the ETL was conjugated to the back aperture of a high-NA objective (NA 1.05, 25×, XLPLN25XWMP2, Olympus) by a pair of achromatic doublets L13 and L14 (L13: f=200 mm, #47-271; L14: f=150 mm, #47-380, Edmund Optics). A phase-only spatial light modulator (HSP1920-1064-HSP8, Meadowlark Optics) was placed at ∼85 mm after L13 and conjugated to the sample plane by the 4f system consisted of L14 and objective lens. The beam diameter entering the objective back aperture is ∼ 11 mm, resulting an excitation NA of ∼0.7. The objective was mounted on a motorized actuator (LTA-HS, Newport) for translation along the optical axis. The excited optical signal was collected by the same objective and directed to the photo-detection unit via dichroic mirror D1 (FF875-Di01-25×36, Semrock). An appropriate band pass filter (FF02-447/60-25, FF03-525/50-25 or FF01-593/46-25, Semrock) along with a short pass filter (FF01-770/SP-25, Semrock) were placed before the photomultiplier tube (PMT) module (H7422, Hamamatsu) to detect the THG/MeOX4, GFP/YFP/GCaMP6s or Texas Red signal. The current signal from PMT was converted to voltage by a transimpedance current amplifier (14 MHz bandwith, ×10^4^ gain, DHCPA-200, Femto).

For 3P imaging, only the stationary beam was selected and the scanning beam was blocked. The AOM was used to control the excitation power by adjusting the driving RF power. Galvanometric scanning mirrors GM2x and GM2y were used for lateral scanning. The excited fluorescence signal was digitized at a sample rate of 120 MHz by a high-speed data acquisition card (vDAQ, Vidrio Technologies). The microscope control software was custom modified based on ScanImage 5.7 ^39^. We synchronized the image acquisition to the laser clock such that each voxel was illuminated by a single or a constant number of excitation pulses. In addition, we only sampled the fluorescence signal over a short time window (∼ 50 ns) for each excitation pulse, while other samples where no signal was presented were rejected to improve the signal-noise-ratio.

### Phase modulation and lock-in detection

DFSS requires a linear phase modulation, equivalent to a frequency shift, of either scanning or stationary beam for lock-in detection (see *Principle of DFSS*). An acousto-optic modulator (AOM) can be used as an optical frequency shifter to generate an accurate linear phase modulation to the weaker scanning beam. However, the AOM efficiency is low for a small frequency shift (e.g. < 1MHz, the repetition rate of excitation laser). Though two cascaded AOMs with slightly different driving frequencies can be used to achieve such a low frequency modulation, the configuration is complicated and efficiency is low. Since the excitation source is a periodic ultrafast laser, we take advantage of this property and achieve low frequency phase modulation with a single AOM. As shown in Fig. S1, after being diffracted by the AOM, the wave function of the scanning beam can be written as:

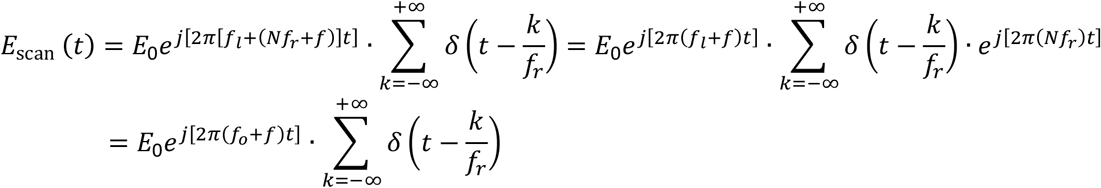

where *f*_*l*_ is the center frequency of excitation laser, *f*_*r*_ is the laser repetition rate and (*Nf*_*r*_ + *f*)is the AOM driving frequency in which *N* is an integer and |*f*| < *f*_*r*_/2. Here, the laser pulse is approximated with a δ-function, because the pulse duration (∼ 60fs) is much shorter than the period of laser cycle (∼ 1µs). Therefore, the frequency shift of (*Nf*_*r*_ + *f*)is equivalent to a linear phase modulation at frequency *f* for a periodic ultrafast laser. This effect is similar to the aliasing phenomenon in signal processing when a sinusoid is sampled at a frequency lower than the Nyquist frequency^40^. In our DFSS-AO-3PM, we chose *f*_*r*_ = 1.002MHz, *f* = 0.4MHz and *N* = 65, resulting in an driving frequency of *Nf*_*r*_ + *f* = 65.5MHz for the AOM.

For signal demodulation, the 3P fluorescence excited by the interference of the stationary and scanning beams was first filtered by a low-pass and band-pass filter (BLP-1.9+ and ZFBP-400K-S+, Mini-Circuit) and then fed into a lock-in amplifier (SR844, Stanford Research Systems). To obtain the reference signal for the lock-in amplifier, the pulse train signal from the oscillator of the Carbide pump laser (∼65.1 MHz), which is the 65th harmonic frequency of the 3P excitation laser, was filtered using a band-pass filter (BBP-60+, Mini-Circuit) and then frequency mixed with the AOM driving signal (65.5 MHz) using a custom-made RF mixer. The output signal from the mixer was filtered by a low-pass filter (BLP-1.9+, Mini-Circuit) and then used as the reference signal (0.4MHz) for the lock-in amplifier. Both the in-phase and quadrature output of the lock-in amplifier were digitized by the data acquisition device and further processed in the control computer.

### Focus sensing and shaping

To measure the PSF, both stationary and scanning beams were focused on the sample. The power of AOM driving signal and the angle of the half wave plate were adjusted such that the intensity ratio of these two beams is ∼10:1. Galvanometric scanner GM2x and GM2y were programmed to park the stationary beam at the target position and scanner GM1 was selected to raster scan the weak probe beam against the strong image beam in a small region (10 ∼ 20 µm). The total laser power was decreased to 70 % of the imaging power during focus sensing to avoid possible photo bleaching or damage. The in-phase (*X*) and quadrature (*Y*) outputs of the lock-in amplifier were recorded, forming the real and imaginary parts of the measured E-field PSF (i.e., *E* = *X* + *iY*, see *Principle of DFSS*). And the corrective phase pattern was the conjugate phase of the Fourier-transform of the E-field PSF. Because the number of pixels of the PSF was much smaller than that of the SLM, zero padding was used before performing the Fourier transform.

To perform optical phase conjugation, the alignment of the SLM and back aperture of the objective needs to be calibrated. The detailed procedure has been reported previously^15^. Briefly, a known phase pattern was first displayed on the SLM and the resulting aberrated PSF was measured using the method described above. The phase of the Fourier transform of the PSF represented the wavefront at the back aperture of the objective lens. Then the affine transformation (rotation, scale and translation) between this wavefront and the phase pattern on the SLM was calibrated. With this information, we could add the affine transformation of the measured wavefront to the SLM and cancel out the aberrations. In conjugate DFSS-AO-3PM system, because the SLM was not placed on the pupil conjugate plane, the diameter and position of laser beam on the SLM would change with the axial and lateral shift of the laser focus. To perform DFSS at any desired position, we conducted a series of calibrations at different focal positions and stored the mapping in a lookup table.

### Conjugate plane calibration

To achieve the optimal FOV for transcranial brain imaging, the SLM was required to be conjugated to the skull plane. The conjugate plane position of the SLM was calibrated by using a fluorescent target (a thin layer of dry rhodamine on a cover glass). First, the ETL was set to 0 diopter position and the fluorescent target was placed at the natural focal plane of the objective using 3P fluorescence as guidance. We recorded the objective position as *z*_1_. Next, a camera was placed before L12 to form a clear image of the SLM via lens L12 and L13. Finally, we translate the objective axially towards the sample using a motorized actuator until the fluorescent target could also be clearly resolved on the camera (i.e., both the fluorescent target and the SLM were imaged on the same plane). The objective position at this step was *z*_2_. The distance between the SLM conjugate plane and objective natural focal plane could be calculated as Δ*z* = *z*_2_ − *z*_1_, which is ∼ 350 µm in our setup. This calibration only need to be performed once and remained unchanged. With this information, we can easily conjugate the SLM to the most aberrated plane in the biological sample. For example, to conjugate the SLM to a mouse skull, we first set ETL to 0 diopter and then located the middle plane of the skull using the intrinsic THG signal. Next, we translated the objective 350 µm towards the sample, which ensured the conjugation of SLM and skull. After that, the position of objective and imaging sample remained unchanged such that the SLM could be always conjugated to the skull. For axial sectioning, the remote focusing module using the ETL was employed.

### Aberration-free remote focusing

The ETL was used to scan the microscope focus axially by controlling the divergence of the excitation light entering the objective. The relationship of the focus shift and ETL control voltage was calibrated using 1-µm-diameter fluorescent beads. Because the infinity-corrected objective lens can only produce diffraction-limited focus for collimated beams, changing the beam divergence will induce large aberration and significantly degrade the imaging performance. To achieve aberration-free remote focusing, a DM was placed on the pupil conjugate plane to compensate for the system aberrations at different focal planes. The aberrations were measured and corrected using a Zernike modal-based sensorless AO algorithm^38^. Briefly, the 3P fluorescence intensity of a fluorescent solution (rhodamine 6G) was chosen as the optimization metric. Nine to eleven different values of each Zernike mode were sequentially applied to the DM and the corresponding fluorescence intensity was Gaussian fitted to find the maximal point. This procedure was repeated for the first 36 Zernike modes (tip, tilt and defocus excluded). Using this method, we calibrated the system aberrations over an axial range of Δ*z* =-300 to 450 µm at an interval of 50 µm, where the natural focal plane of the objective was defined as Δ*z* =0 (i.e., the diopter of ETL was set to 0). For other focal positions, the corrective patterns were linearly interpolated from two nearby calibrated positions. The calibrations for the system aberrations at different focal planes were stored in a lookup table to be used as system AO correction for the conjugate AO imaging over large volume or multiplane imaging.

### DFSS using calcium indicator

Because the fluorescence of the calcium indicator fluctuates, DFSS cannot be directly applied to calcium imaging. To solve this problem, we normalized the outputs of lock-in amplifier (AC term of the 3P signal excited by the interference of scanning and stationary beams) to the fluorescence signal intensity (DC term) during focus sensing (Fig. S11). Briefly, the amplified signal from the PMT was sampled by the data acquisition device and low pass filtered at 10 Hz to extract the fluorescence fluctuation induced by calcium transients. Next, the outputs from the lock-in amplifier were normalized to this value at each scanning coordinate, i.e.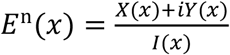, where *X*(*x*)and *Y*(*x*) were the in-phase and quadrature outputs of the lock-in amplifier and *I*(*x*)was the low pass filtered fluorescence intensity.

### Pupil DFSS-AO-3PM system

The optical layout of pupil AO system is similar to the conjugate AO system with only a few modifications. The SLM was moved from the sample-conjugate plane to the pupil conjugate plane and replaced the original DM. The ETL was removed and the actuator carrying the objective was employed for axial sectioning. Lens L5-L8 and L13 were changed accordingly to match the apertures of the SLM, scanning mirrors and objective (L5: f=75, #45-805; L6: f=150 mm, #47-380; L7: f=200 mm, #47-271; L8: f=75 mm, #45-805; L13: f=150 mm, #47-380, Edmund Optics). The final beam diameter is ∼ 15 mm at the objective back aperture, resulting in an excitation NA of 1.05.

#### System aberration correction

The system aberration for the pupil DFSS-AO-3PM system was measured and corrected using DFSS. Briefly, a flat pattern was projected on the SLM and fluorescent beads were used as the imaging target. We parked the strong beam on a fluorescent bead and scanned the weak beam to measure the E-field PSF. Then the phase of the Fourier transform of the PSF was applied to the SLM to correct the system aberration.

### Animal preparation

All the animal experiments were conducted on adult mice (> 2 months old). Five transgenic mouse line: Thy1-YFP (Tg(Thy1-YFP)HJrs/J)^41^, Thy1-GFP-M, Cx3Cr1-GFP (B6.129P2(Cg)-Cx3cr1^tm1Litt^/J)^42^, CCK-GCaMP6s and APP-PS1/Cx3Cr1-GFP mice were used. APP-PS1/Cx3Cr1-GFP mice were obtained by crossing APP-PS1 (Tg(APPswe,PSEN1dE9)85Dbo) mice with Cx3Cr1-GFP mice. CCK-GCaMP6s mice were generated by crossing CCK-ires-Cre (Cck^tm1.1(cre)Zjh^/J) mice with GCaMP6s-Dio (Ai162(TIT2L-GC6s-ICL-tTA2)-D) mice. All animal procedures were conducted in accordance with the Guidelines of the Animal Care Facility of the Hong Kong University of Science and Technology (HKUST) and have been approved by the Animal Ethics Committee at HKUST.

#### Intact skull preparation

Before surgery, mice were anesthetized by intraperitoneal (i.p.) injection of ketamine/xylazine mixture. A midline scalp incision was performed to expose the skull and the connective tissue attached to the skull was gently removed. Then a custom-designed head plate with a circular center hole was glued to the exposed skull for head fixation during *in vivo* imaging. Dental cement was then applied to fill the gap between the head plate and skull and allowed to dry and harden. Finally, the exposed skull was protected by a biocompatible sealant (Kwik-Cast, World Precision Instruments), which can be peeled off before the imaging experiment.

#### Thinned skull window

The surgical procedures for skull exposure and head plate installation were same as preparation of the intact skull. The skull thinning procedures is slightly modified from a previous protocol^43^. Briefly, a 0.5-mm carbon steel burr attached to a high-speed drill was used to thin a small circular region (diameter: 2.0-2.5 mm) centered at 3 mm posterior and lateral to the bregma point. After the majority of the middle spongy bone was removed, a micro surgical blade (no. 6961, Surgistar) was used to thin the skull manually to ∼50 µm thickness, which can be accurately measured using the THG signal of skull bone.

#### Open skull window

Two types of open skull window were used in this study. For the imaging of deep cortex, the cranial window was prepared following a previous protocol ^31^. A craniotomy (4 × 4 mm^2^) over the somatosensory cortex was made using a high-speed drill, leaving the dura intact. A sterile coverslip (4 × 4 mm^2^, 0.2-mm thick) was placed on the exposed dura and glued to the skull using cyanoacrylate adhesive. Then the custom-designed head plate was attached to the skull with dental cement. For the imaging of hippocampus, a whole-brain glass window was implanted according to a previous published approach^32^. Briefly, a custom-made curved glass with a trapezoidal shape (9 × 8 mm^2^, 0.2-mm thick, 10-mm radius curvature) that matched the curvature of mouse brain was used to replace the mouse skull with the dura intact. Then a curved head mount (12-mm radius curvature) was attached to the skull with dental cement.

### *In vivo* imaging

For morphological imaging, the mouse was anesthetized by an intraperitoneal (i.p.) injection of ketamine/xylazine mixture or by inhalation of isoflurane (1-2% in oxygen). For the imaging of vasculature, mice received a retro-orbital intravenous injection of Texas Red (5 mg of dextran conjugate dissolved in 100μl of sterile saline, 70 kDa molecular weight, Thermo Fisher Scientific). For imaging amyloid plaques, the APP-PS1/Cx3Cr1-GFP mice were i.p. injected with MeO-X04 (5.0 mg/kg, 10% DMSO, 90% PBS) 2 hours before imaging. For functional calcium imaging, the mice were under light anesthesia (< 0.5% isoflurane) with their body confined in a tube to reduce motion. During *in vivo* imaging experiments, the mice were mounted on a head-holding stage with angle adjusters (NARISHIGE, MAG-2) and the mouse skull was aligned to be perpendicular to the objective axis by using THG signal of skull as guidance. The detailed imaging parameters including excitation power, pixel dwell time, and DFSS time are summarized in Table. S1.

### Multiplane calcium imaging

Taking advantage of the large corrective volume and quick tuning of focal plane enabled by the conjugate AO and remote focusing technique, we conducted near-simultaneous multiplane calcium imaging. We first measured and corrected the sample-induced aberrations using DFSS and recovered optimal imaging performance over a large volume. We then picked three ROIs and sequentially scan the three ROIs at a frame rate of 10.7Hz using ETL, resulting a volume rate of 3.6 Hz. The DM was synchronized with the ETL to compensate for the divergence-induced system aberrations, which were calibrated before the imaging experiment and stored in a look-up table.

### AO assisted laser microsurgery

To conduct laser-mediated microsurgery, the aberration induced by the biological sample was first corrected using DFSS. Then, the 3P excitation laser was increased to ∼110 mW post objective and focused on the target point for ∼10 s. The injured region was clearly indicated by the new fluorescence signal produced by laser ablation process ^44^.

### Image analysis

The images were processed with Matlab or ImageJ ^45^. To remove the inter-frame motion artefacts, images were registered using the “TurboReg” or “StackReg” plugin ^46^ in ImageJ. Some images were processed with the “Smooth” function in ImageJ, which replaced the value of each pixel with its 3×3 neighboring pixels, or with “Gaussian Blur” function with sigma of 0.15-0.2 µm. The image pairs with system and full AO correction were acquired and processed using the same parameters. For the “system AO” images where the signal intensity was too weak, a linear digital enhancement as indicated in the figures was applied for better visualization.

## Acknowledgements

This work was supported by the Hong Kong Research Grants Council through grants 16103215, 16148816, 16102518, 16102920, T13-607/12R, T13-706/11-1, T13-605/18W, C6002-17GF, C6001-19E, N_HKUST603/19 and the Innovation and Technology Commission (ITCPD/17-9), and the Area of Excellence Scheme of the University Grants Committee (AoE/M-604/16, AOE/M-09/12) and the Hong Kong University of Science & Technology (HKUST) through grant RPC10EG33, and the National Key R&D Program of China (2018YFE0203600). We thank Jufang He, Md Monir Hossian and Mengying Chen from City University of Hong Kong for providing the CCK-GCaMP6s transgenic mice and preparing the open skull window.

## Author contributions

Z.Q. and J.Y.Q. conceived of the research idea. Z.Q. built the AO-3P imaging system and created the control software. Z.Q., Z.S. and C.C. designed and conducted the experiments and data analysis. Z.S. carried out the surgery with the assistance of C.C., Z.Q., W.W. and J.L.. N.Y.I. and J.Y.Q. supervised the project. C.C. and Z.Q. took the lead in writing the manuscript with inputs from all other authors.

## Materials & Correspondence

All data needed to evaluate the conclusions in the paper are present in the paper and/or the Supplementary Materials. Additional data related to this paper may be requested from J.Y.Q.

